# Targeting editing of tomato *SPEECHLESS* cis-regulatory regions generates plants with altered stomatal density in response to changing climate conditions

**DOI:** 10.1101/2023.11.02.564550

**Authors:** Ido Nir, Alanta Budrys, N. Katherine Smoot, Joel Erberich, Dominique C. Bergmann

## Abstract

Flexible developmental programs enable plants to customize their organ size and cellular composition. In leaves of eudicots, the stomatal lineage produces two essential cell types, stomata and pavement cells, but the total numbers and ratio of these cell types can vary. Central to this flexibility is the stomatal lineage initiating transcription factor, SPEECHLESS (SPCH). Here we show, by multiplex CRISPR/Cas9 editing of *SlSPCH cis-*regulatory sequences in tomato, that we can identify variants with altered stomatal development responses to light and temperature cues. Analysis of tomato leaf development across different conditions, aided by newly-created tools for live-cell imaging and translational reporters of SlSPCH and its paralogues SlMUTE and SlFAMA, revealed the series of cellular events that lead to the environmental change-driven responses in leaf form. Plants bearing the novel SlSPCH variants generated in this study are powerful resources for fundamental and applied studies of tomato resilience in response to climate change.

**Significance statement:** Plants can change their shape, size and cellular composition in response to environmental cues. Here, by precise gene editing of a core stomatal development regulator gene in tomato, we generate new alleles with enhanced or dampened responses to light and temperature cues. Combined with live imaging of development, we show the genetic and cellular pathways that contribute to customization of the leaf epidermis, and how this could lead to better climate-adapted varieties.

## Introduction

In response to changing climates, plants adjust their physiology and growth. Clear examples of plant responses are the activity and production of stomata. Stomata are cellular valves in the epidermal surfaces of aerial organs; a pair of stomatal guard cells (GCs) flank a pore whose aperture they modulate through turgor-driven cell swelling. Stomata opening and closing responses occur within minutes of a change in light, water availability, carbon dioxide concentration ([CO_2_]) or temperature, and many of the signal transduction pathways for perception and response have been described in detail (reviewed in (1)). Over longer timescales, stomatal responses to environmental change are detected as changes in stomatal density (SD, stomata/unit area) or stomata index (SI, stomata/total epidermal cell number). Because stomata are preserved in fossils and herbaria, these developmental responses have been used as evidence of past climates (2) (3). Changes to SI and SD can also occur during the development of individual leaves of plants subjected to changes in light (4, 5) or temperature (6–8).

Genes responsible for stomatal production and pattern have been identified, typically first in *Arabidopsis thaliana*, with subsequent studies showing that a core set of transcription factors, receptors and signaling peptides are broadly used among plants (e.g. (9–16)). Among angiosperms, the plants that comprise the majority of natural and cultivated ecosystems, the transcription factors SPEECHLESS (SPCH), MUTE and FAMA play key roles in the specification of stomatal precursors and in the differentiation of the stomatal guard cells and subsidiary cells (reviewed in (17, 18)). EPIDERMAL PATTERNING FACTORS (EPFs) can promote or repress stomatal production and are attractive targets for genetic engineering. Broad overexpression of EPFs can modulate stomatal production and this has measurable effects on important agronomic traits like water use efficiency (WUE) (15, 19).

The stomatal lineage-initiating factor SPCH appears at the nexus of environmental and developmental pattern and, as might be predicted by this position, is subject to extensive regulation at the transcriptional and post-translational levels. Direct regulation of *SPCH* transcription in response to warm temperature is mediated by PHYTOCHROME INTERACTING FACTOR4 (PIF4) (6), and SPCH protein is phosphorylated by MAPK, GSK3 and CDK kinases, (20–22). There is also indirect evidence that SPCH transcript levels and/or protein stability changes when stimuli like light intensity or auxin availability change (23–26).

Advances in precision genome editing have enabled new ways of modulating gene activities, with the potential to also reveal endogenous regulatory mechanisms. Multiplex CRISPR/cas9 mediated editing of tomato gene cis-regulatory regions, for example, revealed enhancers responsible for architecture and fruit traits that mimic those bred into specialized commercial tomato lines, as well as generating novel fruit morphologies (27). Given these technical innovations and the placement of SPCH in stomatal and leaf networks. *SPCH* emerged as an ideal candidate for a CRISPR/CAS9-enabled cis-regulatory region dissection for elements that could drive responsiveness to environmental inputs.

Here we show that *SlSPCH* cis-regulatory alleles can exhibit unique and differential “stomatal set points”, as well as responsiveness to environmental change. To characterize the stomatal response to environmental change at a cellular level in tomato, we created translational reporters of SlSPCH, SlMUTE and SlFAMA, and used a genetically encoded plasma membrane reporter to track cell division patterns in the leaf epidermis over time. Together these new genetic tools revealed that tomato adjusts stomatal production by enabling cells that would typically become stomata to divert into a non-stomatal fate. This adds to fundamental knowledge of plant response and generates potentially useful lines for agronomic use in future climates.

## Results

### Stomatal lineage establishment, expansion and maturation is marked by expression of bHLH transcription factors SlSPCH, SlMUTE and SlFAMA in tomato

Tomato orthologues of the stomatal lineage cell-type specific bHLH transcription factors SPCH, MUTE and FAMA (SlSPCH, SlMUTE and SlFAMA) can complement loss of function mutations in their Arabidopsis mutant counterparts when expressed in Arabidopsis (28), but the expression pattern and role of these tomato genes in their native context has not been described. Knowing these genes’ distribution in tomato is important because, although the epidermal cell type distributions in Arabidopsis and tomato leaves look superficially the same, recent live-cell imaging studies that tracked stomatal lineages over several days identified different modes of cell division in these two plants lead to each species’ final distribution of stomata (Figure 1a) (29).

**Figure 1.**
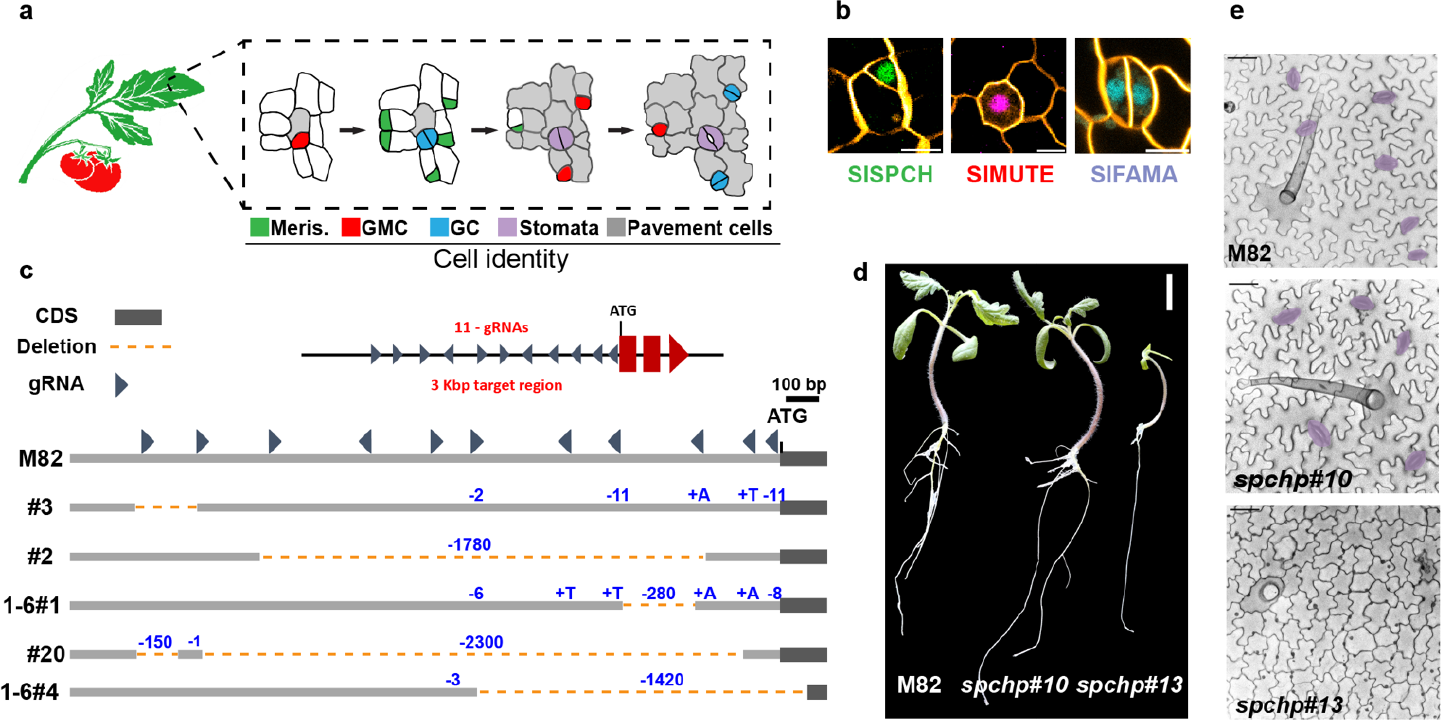
Overall progression of stomatal development in tomato and generation of *SlSPCH* cis-regulatory mutations. (a) Schematic of stomatal development in tomato leaves based on observations of divisions and expression of reporters (b) Confocal images showing cell-type specific expression of stomatal reporters, larger fields of view are in Figure S1a-d. (c) Diagram of *SlSPCH* cis-regulatory mutagenesis scheme and resultant deletions. (d) Seedling morphology and (e) cotyledon epidermis phenotypes in wildtype M82, *SlSPCH* cis-regulatory deletion line 1-6#10 and *SlSPCH* coding region deletion line #13. Stomata are false-colored purple in (e). Scale bars in (b) 10 μm, in (d) 20 mm, and in (e) 50 μm. *See also Figure S1*

We designed native promoter-driven translational reporters and created stable transgenics in tomato cultivar M82 (details in methods). Compared to similar of translational reporters in Arabidopsis, SlSPCHpro::NeonGreen-SlSPCH, SlMUTEpro::mScarlet-SlMUTE, and SlFAMApro::mTurquoise-SlFAMA, were found in roughly the same cell types and patterns (Figure 1b and S1a-d). In newly emerging cotyledons, 0 day after germination (dag) SlSPCH is in many epidermal cells (Figure S1a), but one day later, becomes restricted to the smaller daughters of asymmetric cell divisions (ACDs, Figure S1b). SlMUTE is expressed in cells with the round morphology of guard mother cells (GMCs) (Figure S1c). SlFAMA is expressed in both young guard cells (GCs) before they form a pore (31/48 stomata) and mature GCs with pores (56/69 stomata) (Figure S1d). Unlike Arabidopsis AtFAMA, however, SlFAMA was not detected in GMCs (0/22 GMCs). Under standard long-day growth conditions of 26°C, 700 μmol m^2^ s^-1^ light intensity, there were no obvious overexpression phenotypes generated by the extra copies of SlSPCH, SlMUTE or SlFAMA provided by the transgenes in genetic backgrounds that retained functional copies of the respective endogenous genes.

### Mutagenesis of the *SlSPCH* cis-regulatory region generates small deletions that display a range of stomatal phenotypes

In other species, SPCH has a primary role in lineage flexibility, thus we chose this gene for targeted mutagenesis of its cis-regulatory region. We expected that the complete loss of SlSPCH activity would eliminate stomata and be lethal, thus we employed a CRISPR/Cas9-based multiplex strategy (27) to make a series of small deletions in the 5’ regulatory region of *SlSPCH*. The nearest protein coding gene lies more than 7.5 Kb upstream of the SlSPCH start site (Figure S1e), but ATAC-seq profiles of open chromatin suggested that for most tomato genes, the region ∼3Kb upstream of the start site is likely to contain key regulatory regions (30), and indeed in this smaller region we find numerous predicted target sites for transcription factors with roles in developmental and environmental regulation (Figure S1e). We generated 11 sgRNAs, as evenly distributed as sequence would allow, and generated a series of *SlSPCH* cis-regulatory alleles that range from 2bp to ∼2.5Kbp (Figure 1c and Supplemental file 1), as assayed by genotyping of T1 plants. Plants with deletions spanning more than ∼100 bp were selected and self-pollinated to obtain homozygous lines for all subsequent experiments.

We chose five deletion lines to test for phenotypic response to altered light or temperature conditions. We selected these lines to optimize coverage of the *SlSPCH* cis-regulatory region and we required that when grown under standard growth chamber or greenhouse conditions, the lines were fertile and their overall size and leaf morphology was not substantially different from M82 (Figure 1d-e). Because CRISPR/Cas9 mutagenesis can lead to a spectrum of mutations, we also recovered some plants where deletions disrupted the *SlSPCH* coding region (line #13, Figure 1d-e and S1e-f). These plants were pale, lacked stomata and arrested as seedlings like *Arabidopsis spch* null mutants (31). *SlSPCH* null plants were not characterized further, but served as confirmation that *SlSPCH* is essential for stomatal lineage initiation.

### SlSPCH cis-regulatory mutants display altered responsiveness to changes in light intensity

As the main organs of photosynthesis, leaves have finely tuned responses to changes in light quality and intensity. Along with increasing their mesophyll and chloroplast production, most plants respond to elevated light with an increase in stomatal index (SI) and in total numbers of stomata per leaf. In Arabidopsis, light-mediated promotion of stomatal development is accompanied by higher levels of SPCH. Dissection of the pathway from light perception to stomatal response involves the core light response factor ELONGATED HYPOCOTYL 5 (HY5) upregulating transcription of the mobile signal STOMAGEN in mesophyll cells. STOMAGEN then moves to the epidermis and suppresses the signaling pathway that normally degrades SPCH, ultimately resulting in higher accumulation of SPCH and more stomata (4). It has been suggested, though not experimentally confirmed, that *SPCH* transcription would also be regulated by light (26).

To establish a baseline light response for tomato line M82 (with intact *SlSPCH*), we grew M82 plants at 26°C and two light intensities, ∼130 μmol-photons m^-2^ s^-1^ and ∼1300 μmol-photons m^-2^ s^-1^. We quantified SI, SD and leaf area (Figure 2a-b and S2a-b) as measures of developmental response. As expected, M82 plants increased their SI and SD in the higher light condition. We then tested our *SlSPCH* cis-regulatory deletion mutants in the same conditions and found a variety of responses (Figure 2a). Lines #2, #3 and 1-6#10 have similar SIs to M82 at low light but exhibit a dampened response to higher light intensity. Line #20 has a lower SI at low light, but is similar to M82 at high light, and therefore has an exaggerated response. Line 1-6#4 exhibits both a lower SI at low light and no response to increase in light intensity. Similar trends are seen when SD is calculated (Figure 2b and S2b) except that 1-6#4 does show an increase in SD, but this measurement is sensitive to changes in overall leaf size which typically changes when plants are grown in high light (Figure S2a). Taken together these phenotypes suggest that alteration of the cis-regulatory region of *SlSPCH* can render stomatal production differentially sensitive to light.

**Figure 2.**
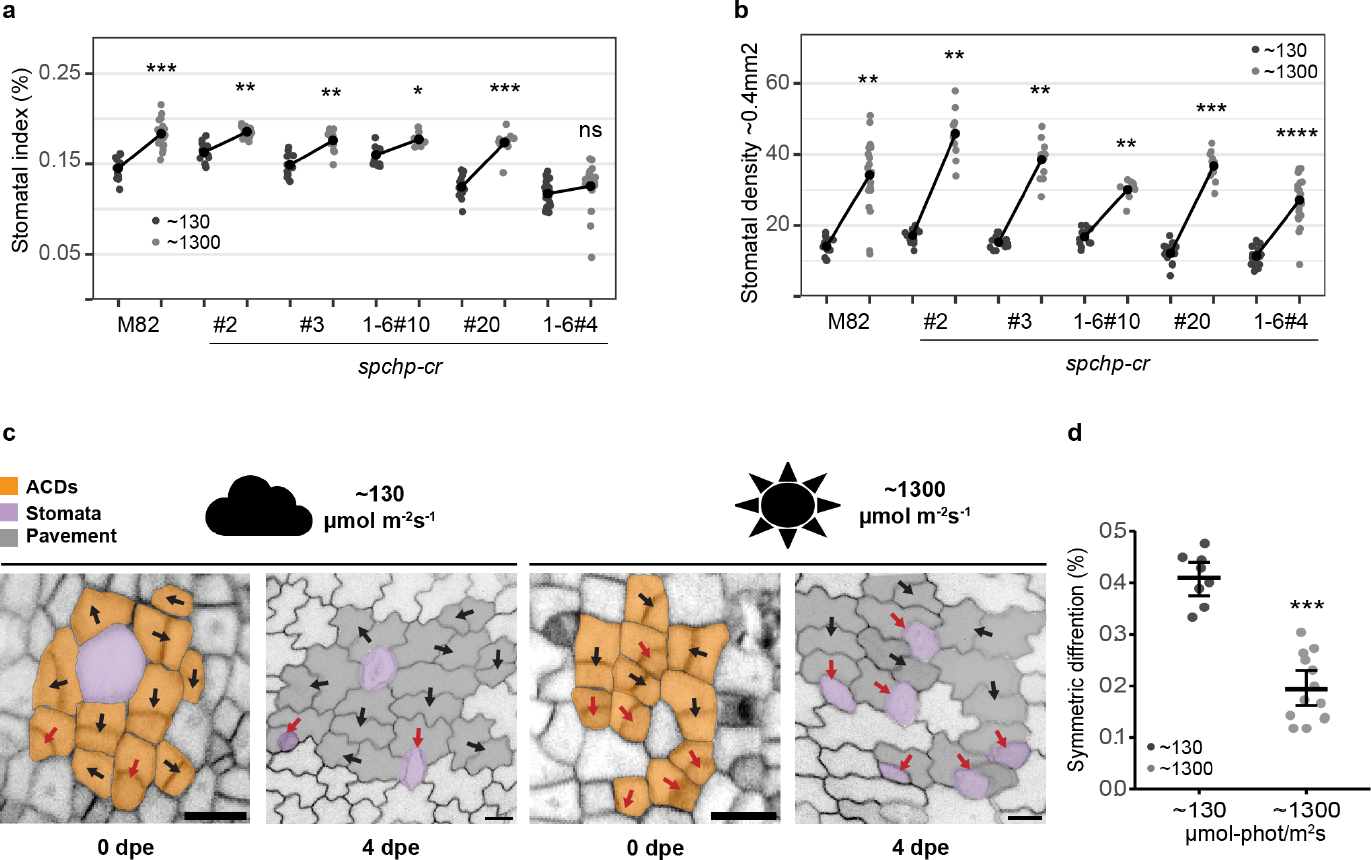
Cellular basis of stomatal response to changes in light intensity and alternation of this response in *SlSPCH* cis-regulatory mutants. (a) Plot of stomatal index (SI) response to low and high light conditions in M82 and cis-regulatory mutants. (b) Plot of stomatal density (SD) response to low and high light conditions in M82 and cis-regulatory mutants.(c) lineage tracing of asymmetrically dividing cells and their fate outcomes from confocal images of M82 cotyledons expressing epidermal plasma membrane reporter *ML1p:RCI2A-NeonGreen*. Red arrows mark ACDs that yield stomata (purple) and black arrows indicate ACDs that lead to two pavement cells (symmetric differentiation). Scale bars represent 20 μm. (d) Plot of shift in the number of ACDs that yield two pavement cells in low and high light. Statistical tests in (a), (b) and (d) are represented as mean ± 95% confidence interval. Bonferroni-corrected p values from Mann-Whitney U test are *P < 0.05; **P < 0.01; ***P < 0.001; ****P < 0.0001. n.s.: P > 0.05, not significant. Sample sizes in (a) and (b) are n = 8-21 0.4mm^2^ fields from 3-5 cotyledons. Sample sizes in (d) are n = 8-14 0.4mm^2^ fields from 3-4 cotyledons. *See also Figure S2*

We wanted to know how tomatoes alter SlSPCH and the stomatal lineage to change SI in response to light. In Arabidopsis, a shift in the relative frequency of amplifying to spacing asymmetric divisions leads to an increase in SI (32). However, in tomato, spacing divisions are nearly non-existent (29), so it was unclear what cellular mechanisms were available for tomato stomatal lineages to alter SI. We therefore imaged plants expressing the epidermal plasma membrane reporter ML1p::RCI2A-NeonGreen (29), every two days to trace asymmetrically dividing precursor cells to their final fate outcome (Figure 2c). From these timecourses, we found that the major mechanism responsible for altering SI was a shift from asymmetric divisions that yield one stoma and one pavement cell to asymmetric divisions that produced two pavement cells (Figure 2d). Thus, tomatoes appear to use a “lineage exit” strategy (29) to lower SI in low light.

### SlSPCH cis-regulatory mutants display altered responsiveness to changes in temperature

Plants also respond to elevated temperature by changing stomatal behavior and production (7, 33). In Arabidopsis, there is evidence for regulation of *SPCH* transcription through repression by the warm-temperature activated PIF4 protein (6). Adapting the temperature-shift protocols in Lau et al., (2018) to account for the different standard temperatures of Arabidopsis and tomato growth (22°C and 26°C respectively), we measured the M82 response to elevated (34°C) temperature, again incorporating lineage tracing to identify the cellular mechanisms responsible for the observed changes in SI and whole leaf attributes. Tomato responses to growth at elevated temperature differed from those in Arabidopsis, most strikingly in the direction of response. In Arabidopsis, higher temperatures suppressed SI, whereas in M82, they led to higher SI and SD (Figure 3a and S2c-d).

**Figure 3.**
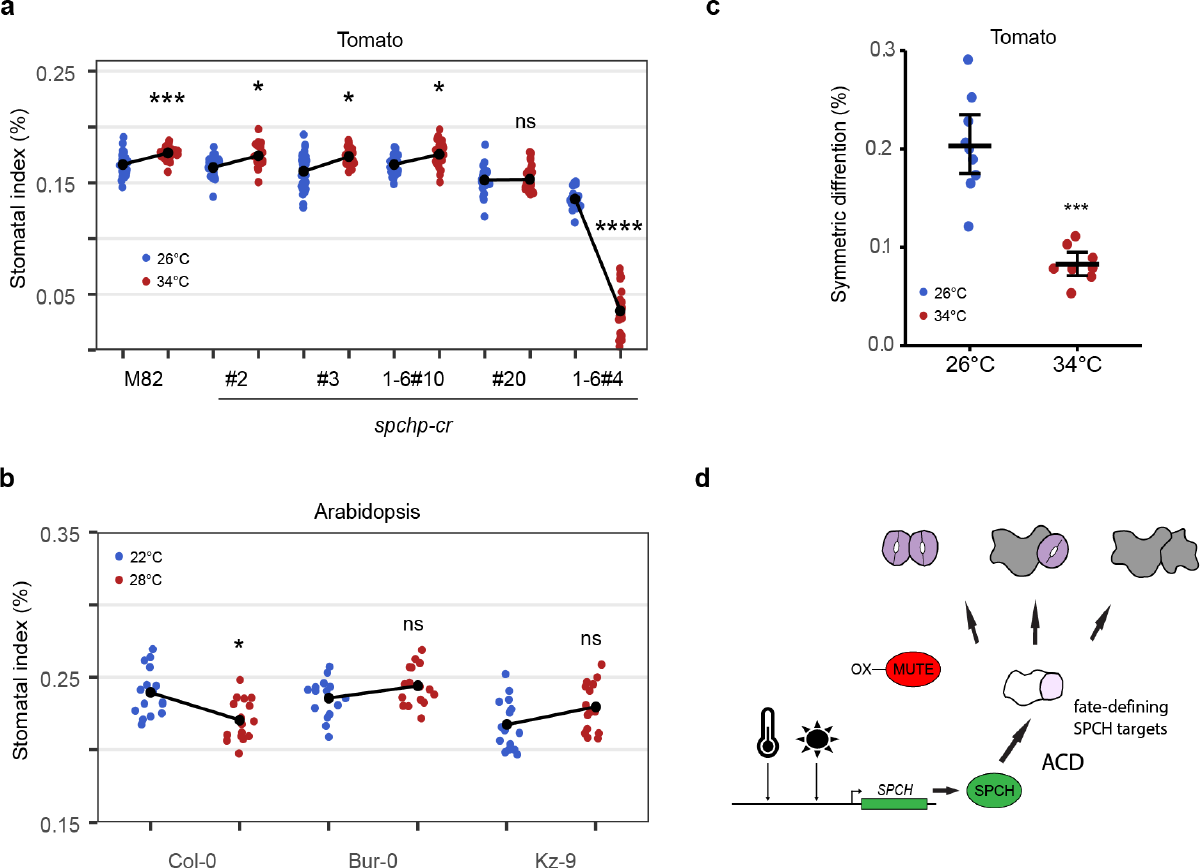
Stomatal response to changes in temperature and alteration of this response in SlSPCH cis-regulatory mutants and among Arabidopsis accessions. Plot of shift in SI of tomato *SlSPCH* cis-regulatory mutants. (b) Plot of SI response of Arabidopsis accessions to change in temperature. (c) Plot of shift in the number of tomato stomatal lineage ACD that produce two pavement cells instead of one stoma and one pavement cell (symmetric differentiation) in plants grown at 26°C and 34°C. (d) Scheme of environmental and genetic influence on the balance between pavement cells and stomata as mediated by SlSPCH and its targets inducing ACDs and then influencing the fates of the daughters. Statistical tests in (a-c) are represented as mean ± 95% confidence interval. Bonferroni-corrected p values from Mann-Whitney U test are *P < 0.05; ***P < 0.001; ****P < 0.0001. n.s.: P > 0.05, not significant. Sample sizes in (a) n = 20-30 0.4mm^2^ fields from 4-6 cotyledons. Sample sizes in (b) n= 15-16 cotyledons each from an individual plant. Sample sizes in (c) are n = 8-9 fields 0.4mm^2^ from 4-6 cotyledons. *See also Figures S2 and S3*

The opposite responses of tomato and Arabidopsis to increased temperature intrigued us. While there are numerous reports of different plants exhibiting stomatal opening and closing responses to changing temperature (34–36), stomatal production or pattern responses are less clear. The Arabidopsis accession used in Lau (2018) was Col-0, a plant from a cool moist climate. Because there are Arabidopsis accessions from diverse climates and with diverse life history traits, we repeated the Arabidopsis temperature-shift experiments with Col-0 and accessions Bur-0 (Ireland) and Kz-9 (Kazakhstan). As before, high temperature results in significantly lower SI in Col-0, but this reduction is not seen in Bur-0 or in Kz-9 (Figure 3b). Additional measurements of SD and leaf area indicate that there is a diversity of responses, and Col-0 alone is not sufficient to fully encompass the “Arabidopsis” response (Figure S2e-f).

Returning to tomato, under the temperature shift regime, the *SlSPCH* cis-regulatory variants again showed diverse responses. SI and SD responses in lines #2, #3 and 1-6#10 to a shift from 26°C to 34°C were slightly dampened relative to those in M82, line *#20* was essentially insensitive, and line *1-6#4* showed a dramatic decrease in SI at 34°C (Figure 3a and S2c-d). The higher temperature resulted in an overall decrease in leaf area in all lines (Figure S2x). Sequencing of line *1-6#4* revealed that a deletion extended into the coding region of *SlSPCH* and thus this response cannot be solely attributed to altering the cis-regulatory region; this allele will be investigated in more detail below (Figure 4). Response to changing temperature could be separated from responses to light in that line #20 was hypersensitive to light, but insensitive to temperature and *1-6#4* only showed a strong change in response to temperature (Figure 2a vs. 3a). Overall, these results show that in addition to the insights CRISPR/Cas9 editing of tomato cis-regulatory regions has generated for morphological diversity (37–39), the approach can reveal sites for environmental inputs.

**Figure 4.**
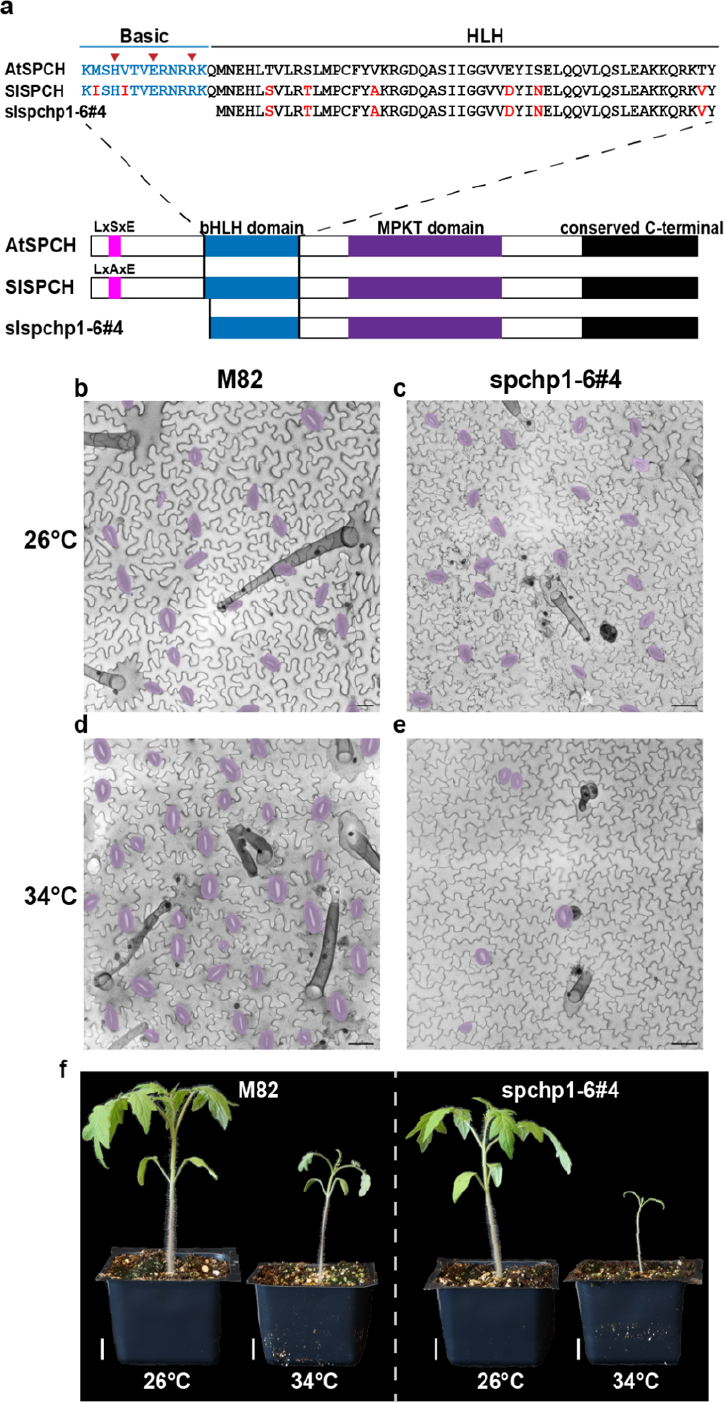
Molecular, cellular and whole-plant characterization of a temperature-sensitive allele of *SlSPCH*. (a) Schematic of SlSPCH protein sequence and predicted effect of 1-6#4 allele. (b-e) Confocal images of cotyledon epidermis in M82 (b.d) and 1-6#4 allele (c, e) at 26C (b-c) and 34C (d-e), stomata false-colored purple. (f) Whole plant images of M82 and SlSPCH 1-6#4 allele at specified temperatures Scale bars in (b-e) 50 μm, (f) 20 mm. See also Figure S4

We used live cell imaging to track how temperature affected stomatal lineage divisions and fates, and found, similar to light responses, that the primary cellular mechanism underlying an increase SI was an increase the proportion of asymmetric precursor divisions producing a stoma and pavement cell (rather than two pavement cells) at higher temperature (Figure 3c). We also took advantage of our SlSPCH, SlMUTE and SlFAMA translational reporters (Figure 1b and S1c and d) to refine how the stomatal lineage responded to changing environments. Due to the broad expression of SlSPCH and the guard cell restricted expression of SlFAMA (Figure S1), we did not gain any additional insights from these reporters. The SlMUTE translational reporter, however, produced a striking phenotype in response to changing temperature (Figure S3). At 34°C, the higher SlMUTE dose provided by the transgene leads to a strong stomatal overproduction and stomatal clustering phenotype (Figure S3a, b and d). We did not see the same phenotype in response to higher light (Figure S3a-c), despite our light shift regime inducing an increase in SI at least as great as our temperature shift regime (Figure 2, 3 and S3). Several models of gene activity are consistent with this phenotype (Figure 3d), and we consider ways in which extra SlMUTE creates a sensitized background that reveals differential behaviors of SlSPCH and asymmetric division programs in the discussion.

### Deletion of coding sequences at the SlSPCH N-terminus generates a temperature sensitive allele of potential agronomic use

The striking and unexpected reduction of SI and SD at high temperature in line *1-6#4* led us to characterize the effects of this mutation on the sequence and function of the resultant SlSPCH protein, as well as on cellular and whole plant phenotypes (Figure 4). The mutation disrupts the N-terminal region of SlSPCH, resulting in a protein that removes the basic residues, including the putative HER motif that mediates DNA binding, but retains the helix-loop-helix domain that is highly conserved among SPCH homologues (Figure 4a). At 26°C, both stomatal production and whole plant phenotypes are indistinguishable from M82, but at 34°C, stomata numbers drop precipitously (Figure 4b-e) and overall growth is stunted (Figure 4f). To assess how a SlSPCH protein missing its N-terminus might behave, we created NeonGreen-tagged reporters and tested expression using a transient expression system in *N. benthamiana*. We infiltrated *N. benthamiana* with a plasmid co-expressing ML1p::RCI2A-mScarletI (control for transformation) and either 35Sp::slspch1-6#4-NeonGreen or 35Sp::SlSPCH-NeonGreen at three temperatures, 22°C, 28°C and 34°C. At all temperatures, 35Sp::SlSPCH-NeonGreen was nuclear, although at higher temperature, fewer cells had detectable signal (Figure S4a, panels 1-3, quantified in S4b). At 22°C, 35Sp::slspch1-6#4 -NeonGreen was also predominantly nuclear, though fewer transformed cells showed expression relative to the full length SlSPCH (Figure S4a, panel 4). At 28°C and 34°C, very little nuclear 35Sp::slspch1-6#4-NeonGreen signal could be detected (Figure S4a, panels 5-6, quantified in S4b). The loss of nuclear localized SlSPCH correlates with the severity of the stomatal phenotype, therefore we conclude that we have generated a temperature sensitive *SlSPCH* allele whose phenotype is linked to its ability to act in the nucleus as a transcription factor. Although we can only speculate about the mechanism underlying the temperature-sensitivity of this *SlSPCH* allele, the ability to create a form of SPCH whose activity can be controlled by temperature opens up new possibilities for testing tomato lines of varying stomatal densities for photosynthetic and water use efficiency phenotypes.

## Discussion

Plants perceive and interpret environmental cues and effect developmental changes. Understanding the signaling, transcriptional and cellular responses that bridge input and output can reveal fundamental mechanisms of information flow and provide materials for creating climate-resilient crops. Here, using CRISPR/Cas9-enabled genome editing of the cis-regulatory region of the stomatal lineage regulatory factor SPCH in tomato, we showed that stomatal production could be made more or less sensitive to light and temperature cues. By combining molecular data on alleles with cellular tracking data, we identified a cell fate switching mechanism that underlies adjustments made in response to light and to temperature. These alleles and new tomato lines bearing reporters for SlSPCH, SlMUTE and SlFAMA will enable finer dissection of stomatal lineage behaviors under many conditions and in other mutant backgrounds. Finally, the identification of a SlSPCH allele whose strong temperature-sensitive activity response appears linked to the subcellular localization of the protein provides both insights into intrinsic regulation of SPCH and a tool for manipulating stomatal production.

Cis-regulatory elements dictate cell-type specific expression and/or can tune expression in response to the environment. We showed that deletions of the 5’ cis-regulatory region of *SlSPCH* result in altered stomatal response to environmental change while leaving overall stomatal patterning (e.g. spacing) intact, suggesting that we identified mainly tuning elements. This outcome was by design, given our initial screen was for CRISPR/Cas9-edited plants with overall wildtype appearance under standard growth conditions. Within the 3Kb region our scRNAs target, there are predicted sites for the light response-mediating transcription factor HY5 (including A-boxes) and two bHLH target sites (E-Boxes) of the type used by the temperature-mediating transcription factor PIF4, and abscisic acid response elements (ABREs). Line #20, which was insensitive to temperature, but hypersensitive to light, disrupts HY5 and two E-boxes (Figure 1c and S1e). Line #10, a relatively small deletion which exhibits a dampened light response, interrupts one of the A-boxes (Figure 1c and S1e). Future experiments may reveal the roles of tomato HY5 and PIF genes (40, 41) in stomatal regulation. It will be particularly interesting to find that HY5 is a direct transcriptional regulator of *SlSPCH* because in Arabidopsis, there is an intermediary between HY5 and regulation of SPCH protein (4). Sequence predictions in the *SlSPCH* upstream region, and alignment with SPCH 5’ regions from other Solanaceae and from Arabidopsis reveal a number of conserved regions (CNS, Figure S1e). Some of the Solanaceae CNS are included in our characterized deletion lines (Figure 1c and S1e) but the CNS conserved between tomato and Arabidopsis (red in Figure S1e) lie further upstream. Generation of additional cis-regulatory alleles that specifically target CNS may be a productive strategy to identify additional regulatory elements, including both those that drive cell-type specific expression and those that affect *SlSPCH* production in response to additional environmental inputs.

The behavior of the SlSPCH1-6#4 allele that eliminates the N-terminal region, including the predicted DNA-binding region of the bHLH domain, challenges our assumptions about which regions of the SlSPCH protein are essential, and for which activities. At 26°C *SlSPCH1-6#4* lines produce slightly fewer stomata than M82 (Figure 3a), but the stomata appear morphologically normal and well-patterned (Figure 4b and c), therefore this version of SlSPCH must be capable of performing the general SlSPCH functions of inducing asymmetric cell divisions and promoting eventual guard cell identity. Although it may seem surprising to ascribe transcriptional regulator activity to a bHLH missing the motif that typically mediates DNA contacts, there are classes of transcription factors that can function without sequence N-terminal of the HLH domains (42), and previous deletion and domain swap experiments showed that AtMUTE, and to a lesser extent, AtSPCH, can function when three putative DNA-binding residues (HER) are replaced (43). In other plants, SPCH is an obligate heterodimer working with INDUCER OF CBF EXPRESSION (ICE1) and paralogous ClassIIIb bHLHs (11, 16, 44). It is possible that as a heterodimer, it is the ICE1 residues that make contact with DNA, and SPCH provides some other essential activities for transcriptional activation (45). Loss of the SlSPCH N-terminal region is also predicted to eliminate a nuclear localization signal, and we see that, at high temperatures, SlSPCH1-6#4 cannot be retained in the nucleus (Figure S4). That SlSPCH1-6#4 is ever nuclear we attribute to its heterodimerization with *SlICE*. We speculate that, given ICE1’s activity promoting gene expression at low temperatures (46, 47), it may be less present or active at warm ones, and low ICE1 would be insufficient to retain the N-terminal deleted SlSPCH.

Ultimately how do the manipulations we made to the *SlSPCH* locus lead to environment-dependent changes in stomatal production? Cell tracking data in M82 suggest that SlSPCH-induced ACDs normally resolve into either a stoma and pavement cell or two pavement cells, and light and temperature modulate SlSPCH production and/or activity to tip this balance (Model in Figure 3d). One hypothesis is a quantitative shift in *SlSPCH* in cells about to undergo ACDs defines whether one or no cells acquire stomatal fate after ACD. We, however, prefer a second explanation that also takes into account that in response to higher temperature, pairs or small clusters of stomata result from the presence of SlMUTEpro::SlMUTE-mScarletI. Broad overexpression of MUTE in Arabidopsis leads to the formation of extra stomata by inducing non-stomatal cells to take on stomatal fate (48). The ectopic stomata produced by the extra copy of *SlSMUTE*, however, are morphologically normal, and the orientation of the GC pairs suggests that they do not originate from extra divisions of GCs (Figure S3, compared to (49)). Therefore, we hypothesize that the extra SlMUTE dosage is revealing more of the range in the post-ACD toggle between pavement cell and GC identity. As diagrammed in Figure 3d, this suggests that tomato ACDs produce a continuum of cell fates. Under conditions optimal for productive photosynthesis and leaf growth, ACDs result in one GC and one pavement cell. The ACDs are induced by SlSPCH, but the resulting choice of fate is dictated by SlSPCH targets (or SlSPCH in complex with a target) and among those targets is *SlMUTE*. Under limiting environmental conditions, SlSPCH levels or activity are insufficient to induce fate-promoting targets, and both products of an ACD become pavement cells. In high temperature conditions, elevated SlSPCH and/or its direct targets would have both endogenous and transgene-derived SlMUTE to induce and this higher expression would push the ACDs to yield two stomata. Alternatively, SlMUTE itself could be temperature responsive and an additional copy of SlMUTE could cause SlMUTE to accumulate to levels high enough to pass a cell fate threshold in both daughters of an ACD.

Nearly all future climate predictions suggest that temperatures and atmospheric [CO_2_] will increase, and severe weather events (drought, flooding) will become more prevalent (50, 51). Stomata, with their roles in capturing atmospheric carbon and regulating plant transpiration, which has both water transport and leaf cooling components, must integrate and prioritize potentially conflicting signals to maintain plant health. While the increase in stomatal production in response to increasing light intensity appears to occur in most plants, the response to increased temperature can vary, as seen comparing tomato and Arabidopsis, and even among Arabidopsis accessions. These diverging responses to warm temperature may reflect different priorities for conserving water and leaf cooling. It is therefore hard to predict, but important to test, the combinatorial effects of predicted future climate conditions on plants. Genetic tools that can alter sensitivity to specific inputs, such as the SlSCPH lines generated in this work, will be instrumental in deciphering complex response and may also be useful to assay the effects of having tomato plants with different stomatal numbers for growth in large-scale or urban agricultures systems of the future.

## Supporting information

Supplemental data: Sequence alignments

## Author Contributions

N designed research, performed research, analyzed data, and wrote the paper; AB and NKS performed research, JE performed research and analyzed data, DCB designed research, analyzed data and wrote the paper.

## References

1. C. B. Engineer, et al., CO2 Sensing and CO2 Regulation of Stomatal Conductance: Advances and Open Questions. Trends Plant Sci. 21, 16–30 (2016).

2. J. C. McElwain, M. Steinthorsdottir, Paleoecology, Ploidy, Paleoatmospheric Composition, and Developmental Biology: A Review of the Multiple Uses of Fossil Stomata. Plant Physiol. 174, 650–664 (2017).

3. P. L. M. Lang, et al., Century-long timelines of herbarium genomes predict plant stomatal response to climate change. bioRxiv, 2022.10.23.513440 (2022).

4. S. Wang, et al., Light regulates stomatal development by modulating paracrine signaling from inner tissues. Nat. Commun. 12, 3403 (2021).

5. M. F. Pompelli, S. C. V. Martins, E. F. Celin, M. C. Ventrella, F. M. Damatta, What is the influence of ordinary epidermal cells and stomata on the leaf plasticity of coffee plants grown under full-sun and shady conditions? Braz. J. Biol. 70, 1083–1088 (2010).

6. O. S. Lau, et al., Direct Control of SPEECHLESS by PIF4 in the High-Temperature Response of Stomatal Development. Curr. Biol. 28, 1273–1280.e3 (2018).

7. Y. Zheng, et al., Effects of experimental warming on stomatal traits in leaves of maize (Zea may L.). Ecol. Evol. 3, 3095–3111 (2013).

8. S. Marek, et al., Stomatal density in Pinus sylvestris as an indicator of temperature rather than CO2 : Evidence from a pan-European transect. Plant Cell Environ. 45, 121–132 (2022).

9. H. Wang, et al., BZU2/ZmMUTE controls symmetrical division of guard mother cell and specifies neighbor cell fate in maize. PLoS Genet. 15, e1008377 (2019).

10. M. T. Raissig, et al., Mobile MUTE specifies subsidiary cells to build physiologically improved grass stomata. Science 355, 1215–1218 (2017).

11. M. T. Raissig, E. Abrash, A. Bettadapur, J. P. Vogel, D. C. Bergmann, Grasses use an alternatively wired bHLH transcription factor network to establish stomatal identity. Proc. Natl. Acad. Sci. U. S. A. 113, 8326–8331 (2016).

12. Z. Jiao, et al., PdEPFL6 reduces stomatal density to improve drought tolerance in poplar. Ind. Crops Prod. 182, 114873 (2022).

13. M. Clemens, et al., VvEPFL9-1 Knock-Out via CRISPR/Cas9 Reduces Stomatal Density in Grapevine. Front. Plant Sci. 13, 878001 (2022).

14. C. C. C. Chater, R. S. Caine, A. J. Fleming, J. E. Gray, Origins and Evolution of Stomatal Development. Plant Phys. 174, 624–638 (2017).

15. R. S. Caine, et al., Rice with reduced stomatal density conserves water and has improved drought tolerance under future climate conditions. New Phytol. 221, 371–384 (2019).

16. C. C. Chater, et al., Origin and function of stomata in the moss Physcomitrella patens. Nat Plants 2, 16179 (2016).

17. K. H. McKown, D. C. Bergmann, Stomatal development in the grasses: lessons from models and crops (and crop models). New Phytol. 227, 1636–1648 (2020).

18. L. R. Lee, D. C. Bergmann, The plant stomatal lineage at a glance. J. Cell Sci. 132 (2019).

19. T. Doheny-Adams, L. Hunt, P. J. Franks, D. J. Beerling, J. E. Gray, Genetic manipulation of stomatal density influences stomatal size, plant growth and tolerance to restricted water supply across a growth carbon dioxide gradient. Philos. Trans. R. Soc. Lond. B Biol. Sci. 367, 547–555 (2012).

20. G. R. Lampard, C. A. Macalister, D. C. Bergmann, Arabidopsis stomatal initiation is controlled by MAPK-mediated regulation of the bHLH SPEECHLESS. Science 322, 1113–1116 (2008).

21. G. E. Gudesblat, et al., SPEECHLESS integrates brassinosteroid and stomata signalling pathways. Nat. Cell Biol. 14, 548–554 (2012).

22. K.-Z. Yang, et al., Phosphorylation of Serine 186 of bHLH Transcription Factor SPEECHLESS Promotes Stomatal Development in Arabidopsis. Mol. Plant 8, 783–795 (2015).

23. D. Samakovli, et al., YODA-HSP90 Module Regulates Phosphorylation-Dependent Inactivation of SPEECHLESS to Control Stomatal Development under Acute Heat Stress in Arabidopsis. Mol. Plant 13, 612–633 (2020).

24. X. Yang, L. Gavya S, Z. Zhou, D. Urano, O. S. Lau, Abscisic acid regulates stomatal production by imprinting a SnRK2 kinase-mediated phosphocode on the master regulator SPEECHLESS. Sci Adv 8, eadd2063 (2022).

25. C. Han, et al., TOR and SnRK1 fine tune SPEECHLESS transcription and protein stability to optimize stomatal development in response to exogenously supplied sugar. New Phytol. 234, 107–121 (2022).

26. C. Klermund, et al., LLM-Domain B-GATA Transcription Factors Promote Stomatal Development Downstream of Light Signaling Pathways in Arabidopsis thaliana Hypocotyls. Plant Cell 28, 646–660 (2016).

27. D. Rodríguez-Leal, Z. H. Lemmon, J. Man, M. E. Bartlett, Z. B. Lippman, Engineering Quantitative Trait Variation for Crop Improvement by Genome Editing. Cell 171, 470–480.e8 (2017).

28. A. Ortega, A. de Marcos, J. Illescas-Miranda, M. Mena, C. Fenoll, The Tomato Genome Encodes SPCH, MUTE, and FAMA Candidates That Can Replace the Endogenous Functions of Their Arabidopsis Orthologs. Front. Plant Sci. 10, 1300 (2019).

29. I. Nir, et al., Evolution of polarity protein BASL and the capacity for stomatal lineage asymmetric divisions. Curr. Biol. 32, 329–337.e5 (2022).

30. A. Hendelman, et al., Conserved pleiotropy of an ancient plant homeobox gene uncovered by cis-regulatory dissection. Cell 184, 1724–1739.e16 (2021).

31. C. A. MacAlister, K. Ohashi-Ito, D. C. Bergmann, Transcription factor control of asymmetric cell divisions that establish the stomatal lineage. Nature 445, 537–540 (2007).

32. A. Vatén, C. L. Soyars, P. T. Tarr, Z. L. Nimchuk, D. C. Bergmann, Modulation of Asymmetric Division Diversity through Cytokinin and SPEECHLESS Regulatory Interactions in the Arabidopsis Stomatal Lineage. Dev. Cell 47, 53–66.e5 (2018).

33. J. Zhu, et al., Effect of simulated warming on leaf functional traits of urban greening plants. BMC Plant Biol. 20, 139 (2020).

34. Y. Li, et al., SlSnRK2.3 interacts with SlSUI1 to modulate high temperature tolerance via Abscisic acid (ABA) controlling stomatal movement in tomato. Plant Sci. 321, 111305 (2022).

35. J. Urban, M. W. Ingwers, M. A. McGuire, R. O. Teskey, Increase in leaf temperature opens stomata and decouples net photosynthesis from stomatal conductance in Pinus taeda and Populus deltoides x nigra. J. Exp. Bot. 68, 1757–1767 (2017).

36. K. Gasparini, et al., The Lanata trichome mutation increases stomatal conductance and reduces leaf temperature in tomato. J. Plant Physiol. 260, 153413 (2021).

37. M. Alonge, et al., Major Impacts of Widespread Structural Variation on Gene Expression and Crop Improvement in Tomato. Cell 182, 145–161.e23 (2020).

38. X. Wang, et al., Dissecting cis-regulatory control of quantitative trait variation in a plant stem cell circuit. Nat Plants 7, 419–427 (2021).

39. C.-T. Kwon, et al., Rapid customization of Solanaceae fruit crops for urban agriculture. Nat. Biotechnol. 38, 182–188 (2020).

40. D. Rosado, et al., Phytochrome Interacting Factors (PIFs) in Solanum lycopersicum: Diversity, Evolutionary History and Expression Profiling during Different Developmental Processes. PLoS One 11, e0165929 (2016).

41. C. Zhang, et al., Pivotal roles of ELONGATED HYPOCOTYL5 in regulation of plant development and fruit metabolism in tomato. Plant Physiol. 189, 527–540 (2022).

42. Y. Hao, X. Zong, P. Ren, Y. Qian, A. Fu, Basic Helix-Loop-Helix (bHLH) Transcription Factors Regulate a Wide Range of Functions in Arabidopsis. Int. J. Mol. Sci. 22 (2021).

43. K. A. Davies, D. C. Bergmann, Functional specialization of stomatal bHLHs through modification of DNA-binding and phosphoregulation potential. Proc. Natl. Acad. Sci. U. S. A. 111, 15585–15590 (2014).

44. M. M. Kanaoka, et al., SCREAM/ICE1 and SCREAM2 specify three cell-state transitional steps leading to arabidopsis stomatal differentiation. Plant Cell 20, 1775–1785 (2008).

45. A. Liu, et al., Cell Fate Programming by Transcription Factors and Epigenetic Machinery in Stomatal Development. bioRxiv, 2023.08.23.554515 (2023).

46. T. Ma, et al., Arabidopsis LFR, a SWI/SNF complex component, interacts with ICE1 and activates ICE1 and CBF3 expression in cold acclimation. Front. Plant Sci. 14, 1097158 (2023).

47. R. Lin, et al., CALMODULIN6 negatively regulates cold tolerance by attenuating ICE1-dependent stress responses in tomato. Plant Physiol. 193, 2105–2121 (2023).

48. L. J. Pillitteri, D. B. Sloan, N. L. Bogenschutz, K. U. Torii, Termination of asymmetric cell division and differentiation of stomata. Nature 445, 501–505 (2007).

49. L. B. Lai, et al., The Arabidopsis R2R3 MYB proteins FOUR LIPS and MYB88 restrict divisions late in the stomatal cell lineage. Plant Cell 17, 2754–2767 (2005).

50. S. Alamos, P. M. Shih, How to engineer the unknown: Advancing a quantitative and predictive understanding of plant and soil biology to address climate change. PLoS Biol. 21, e3002190 (2023).

51. M. Haworth, G. Marino, F. Loreto, M. Centritto, Integrating stomatal physiology and morphology: evolution of stomatal control and development of future crops. Oecologia 197, 867–883 (2021).

