## Supplemental figures and methods for "Targeting editing of tomato *SPEECHLESS* cis-regulatory regions generates plants with altered stomatal density in response to changing climate conditions"

[3] Current Address, Institute of Plant Sciences, ARO, Volcani Center, HaMaccabbim Road 68, POB 15159, Rishon LeZion 7505101, Israel.

[4] Current Address, Department of Biology, New York University, 24 Waverly Pl, New York, NY, 10003, USA

[5] Current Address, Plant and Microbial Biology, UC Berkeley, Berkeley, CA 94720, USA

### Contains:

### Methods

### Supplemental Figures

**Figure S1** Extended stomatal gene reporter expression and protein alignments

**Figure S2** Additional SI, SD and leaf area measurements of tomato and Arabidopsis variants in response to changing environmental conditions

**Figure S3** Tomato lines bearing a SlMUTE translational reporter display a hypersensitive response to increased temperature, but not light

**Figure S4** Nuclear localization changes in response to high temperature in *SlSPSCH* 1-6#4.

### Supplemental Tables

**Table S1** List of primers used for constructing reporters and for genotyping

**Table S2** List of SlSPCH promoter sgRNAs used for CRISPR/Cas9 mutagenesis and PCR primers used for genotyping

### Methods

Plant growth: Tomato plants (*Solanum lycopersicum* cv. M82) were grown in a Percival chamber model CU22L, or Percival growth room, model AR-1015L3 set to a photoperiod of 16 hr light/8 hr dark, light intensity of approximately 250 µmol m2 s-1, and 26 °C or 34 °C. For propagation, plants were grown in a greenhouse under natural day length conditions, at 700–1200 µmol m2 s-1 and 18 – 29 °C. For light experiments the Percival AR-1015L3 growth room light intensity set to 130 µmol $m^2$ $s^{-1}$ for low light experiment and 1300 µmol $m^2$ $s^{-1}$ for high light experiment.

Tomato DNA constructs and plant transformation: For tomato SlSPCH, SlMUTE, and SlFAMA marker lines, the promoters and CDS (primers used for cloning are in Table S1) were cloned into a level 0 MoClo part using the Golden Gate cloning system (1) and then fused to NeonGreen, mScarlet and mTurquoise respectively with a NOS terminator to form a level 1 construct. Each level 1 was transferred to a level 2, together with a kanamycin resistance cassette. The constructs were sub-cloned into the pAGM4723 binary vector and were introduced into *Agrobacterium tumefaciens* strain GV3101 by electroporation. The constructs were transferred to M82 cotyledons, using transformation and regeneration methods described by (2). Kanamycin-resistant T0 plants were grown and at least four independent transgenic lines were selected and self-pollinated to generate homozygous transgenic lines.

Tomato SPCH promoter CRISPR/Cas9 mutagenesis, plant transformation, and selection of mutant alleles 11 single-guide RNAs (sgRNAs) spanning 3000 bases upstream of the TSS of SlSPCH were designed using the CRISPR-P tool (3). Primers used for sgRNAs and subsequent genotyping are in Table S2. The gRNAs and promoter were assembled using the Golden Gate cloning system as described in (1). The final binary vector including zCas9, the gRNAs and NPTII, assembled in pAGM4723, was introduced into *Agrobacterium tumefaciens* strain GV3101 by electroporation. The construct was transferred into M82 cotyledons using transformation and regeneration methods described by (2). T0 transgenic plants resistant to Kanamycin were grown and independent lines were selected and self-pollinated to generate homozygous lines. For genotyping of the transgenic lines, genomic DNA was extracted, and each plant was genotyped by PCR for the presence of the zCas9. The positive lines for the zCas9 were further genotyped for mutations in SlSPCH promoter (Solyc03g007410) using a forward primer 3.1Kb upstream to the ATG and a reverse primer 155 bp downstream to the ATG, these pair primers cover the 11 gRNAs

Microscopy, image analysis and processing The fluorescence imaging experiments on the tomato transgenic plants were performed on a Leica SP5, SP8 and Stellaris confocal microscope with HyD detectors using 25X NA0.95 and 40x NA1.1 water objective with image size 1024*1024 and digital zoom from 1x to 2x. To quantify the stomatal index of the tomato SPCH promoter mutants stomatal, 14 dpg cotyledons were imaged and the fraction of stomata from total cells in leaf epidermis was computed from regions of approximately 0.4 mm2. Stomatal index was calculated as- S/(S+E), S- number of stomata, E- number of epidermis cells. Still or time-course images of ML1p::RCI2A-NeonGreen, propidium iodide (PI), and FM4-64 fluorescence in tomato were obtained from a Leica Stellaris confocal microscope with HyD detectors using 25x water objective with image size 1024*1024 and digital zoom from 1x to 2x. Time-course experiments on tomato cotyledons were done as described by (4). All raw fluorescence image Z-stacks were projected with STD Slices in FIJI.

Transient expression in *Nicotiana benthamiana*: *N. benthamiana* plants were grown for six weeks in a phytotron greenhouse under natural day length conditions, at 700–1200 mmol m-2 s-1, with daytime temperature of 22°C, then moved to 22°C, 28°C, and 34°C for Agrobacterium infiltration and subsequent analysis of protein expression as indicated. Plants were at least 6 weeks old at time of experiment. For transient expression, dual expression constructs with expression control ML1pro::RCI2A-mScarlet-I and either 35Spro:SlSPCH-NeonGreen or 35Spro:slspch1-6#4-NeonGreen were cloned using the Golden Gate system (1) into the binary vector pAGM4723 with rbcS terminator for SPCH variants, and NOS terminator for control. The binary vectors were introduced into *Agrobacterium tumefaciens* strain GV3101 and infiltrated into mature tobacco leaves as described in (5).

The analysis of stomatal density and index responses to changes in temperature in Arabidopsis of different accessions, followed growth condition and temperature shift protocols in (6). Seeds for accessions Col-0 (CS22625), Bur-0 (CS77833) and Kz-9 (CS22607) were obtained from the ABRC, Ohio State University, USA. Tissues were cleared by placing 7:1 Ethanol:Acetic acid for 3 days, rinsed in 1M Potassium hydroxide for 30 minutes, then placed in water before mounted with Hoyer's solution. DIC images of the abaxial surface were collected at 20X using a Leica-DFC9000GTC-VSC11962 with a 1.623 $mm^2$ field of view and 50 µm z-step size. Stomata for density were counted across the entire field of view whereas stomata and other epidermal cells for index were counted on a 0.370 $mm^2$

subset placed on a non-vein section of the lamina. Cells were counted using ImageJ with the CellCounter plugin. Results were plotted using RStudio with the "tidyverse" suite of packages.

Gene identifiers: *Solanum lycopersicum* v.M82: SlSPCH (Solyc03g007410); SlMUTE (Solyc1g080050); SlFAMA (Solyc05g05366010) *Arabidopsis thaliana:* SPCH (At5g53210), MUTE (At3g06120) FAMA (At3g24140),

Quantification and statistical analyses

All statistical analyses in this manuscript were performed in RStudio (R Development Core Team., 2020). Statistical parameters for each analysis are indicated in the figure legends. For significance testing, unpaired Mann-Whitney *U* tests were conducted with the wilcox_test function from the rstatix package (7). Bonferroni corrections were performed when more than 2 pair-wise comparisons were conducted, and Bonferroni corrected p-values are indicated in all figures where applicable.


1. E. Weber, R. Gruetzner, S. Werner, C. Engler, S. Marillonnet, Plos One (2011).

2. S. McCormick, "Transformation of tomato with Agrobacterium tumefaciens" in *Plant Tissue Culture Manual*, (Springer, 1991), pp. 311–319.

3. Y. Lei, *et al.*, CRISPR-P: a web tool for synthetic single-guide RNA design of CRISPR-system in plants. *Mol. Plant* **7**, 1494–1496 (2014).

4. I. Nir, *et al.*, Evolution of polarity protein BASL and the capacity for stomatal lineage asymmetric divisions. *Curr. Biol.* **32**, 329–337.e5 (2022).

5. Y. Zhang, P. Wang, W. Shao, J.-K. Zhu, J. Dong, The BASL polarity protein controls a MAPK signaling feedback loop in asymmetric cell division. *Dev. Cell* **33**, 136–149 (2015).

6. O. S. Lau, *et al.*, Direct Control of SPEECHLESS by PIF4 in the High-Temperature Response of Stomatal Development. *Curr. Biol.* **28**, 1273–1280.e3 (2018).

7. A. Kassambara, rstatix: Pipe-friendly framework for basic statistical tests. R package version 0.7. 0 (2021).

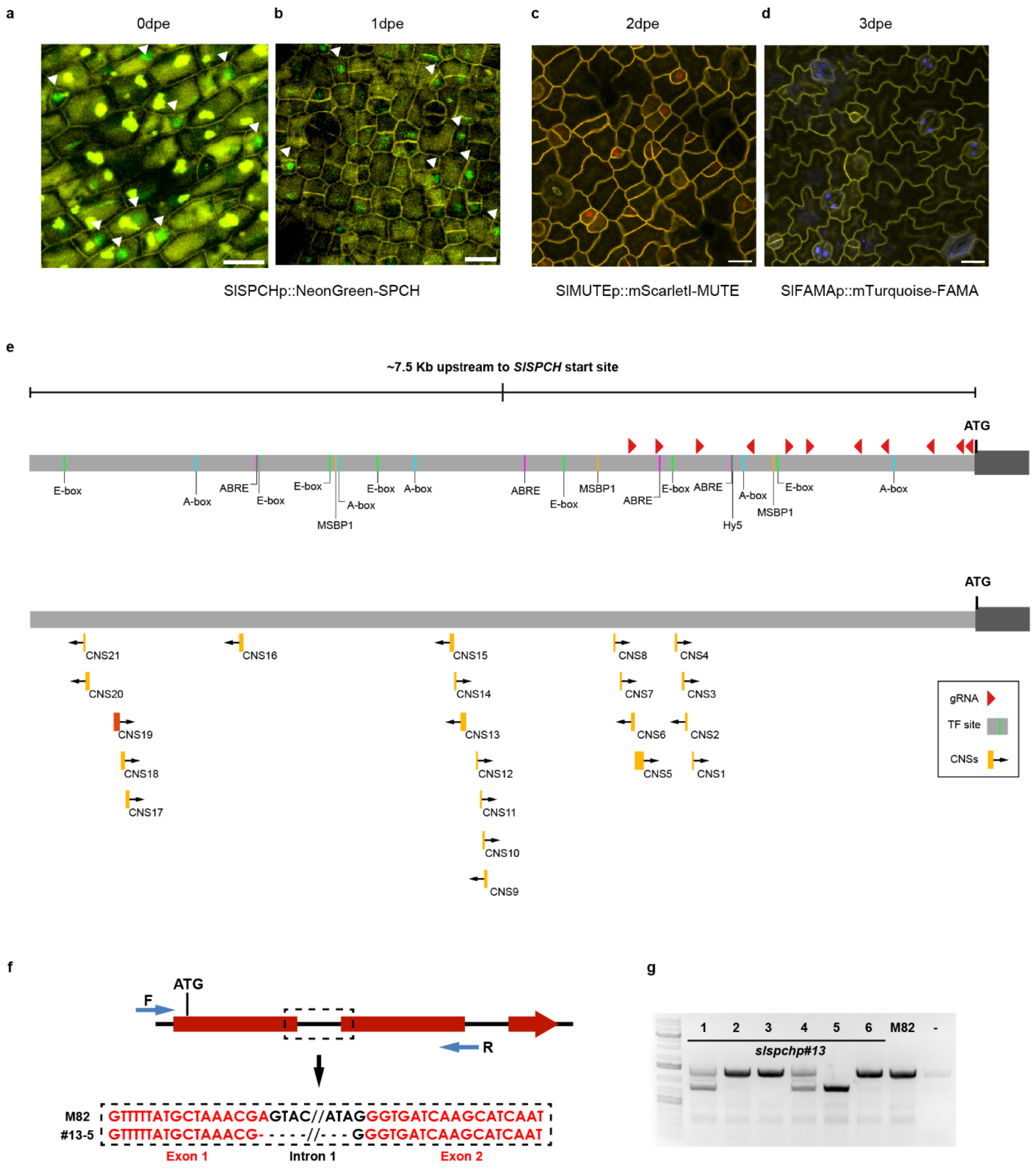


**Figure S1 Extended stomatal gene reporter expression and protein alignments (related to Figure 1)**
(a-d) Confocal images of stomatal lineage reporters on days of peak expression in M82 cotyledons. Scale bars represent 20 µm. (e) Annotation of SlSPCH 5' cis-regulatory region with gRNAs (red triangles), predicted transcription factor (TF) binding sites, and Conserved Non-Coding Sequences (CNS) found between *S. lycopersicum* and other plant species. Orange CNSs are conserved only within the Solanaceae family, and the red CNS (CNS19) is conserved among eudicots including Arabidopsis. (f) Description of mutation in SlSPCH line #13 (phenotype in Figure 1). This putative *SlSPCH* null allele (frameshift in coding region) has a deletion (dashed square) more than 300 bp away from gRNAs. The black arrow points to the exon (red) and intron (black) sequences that are deleted in line #13. (g) PCR genotyping of offspring from plant heterozygous for the #13 allele showing segregation: plants 1 and 4 are heterozygotes, 5 is a homozygous mutant and plants 2, 3 and 6 are wild-type at the *SlSPCH* locus.

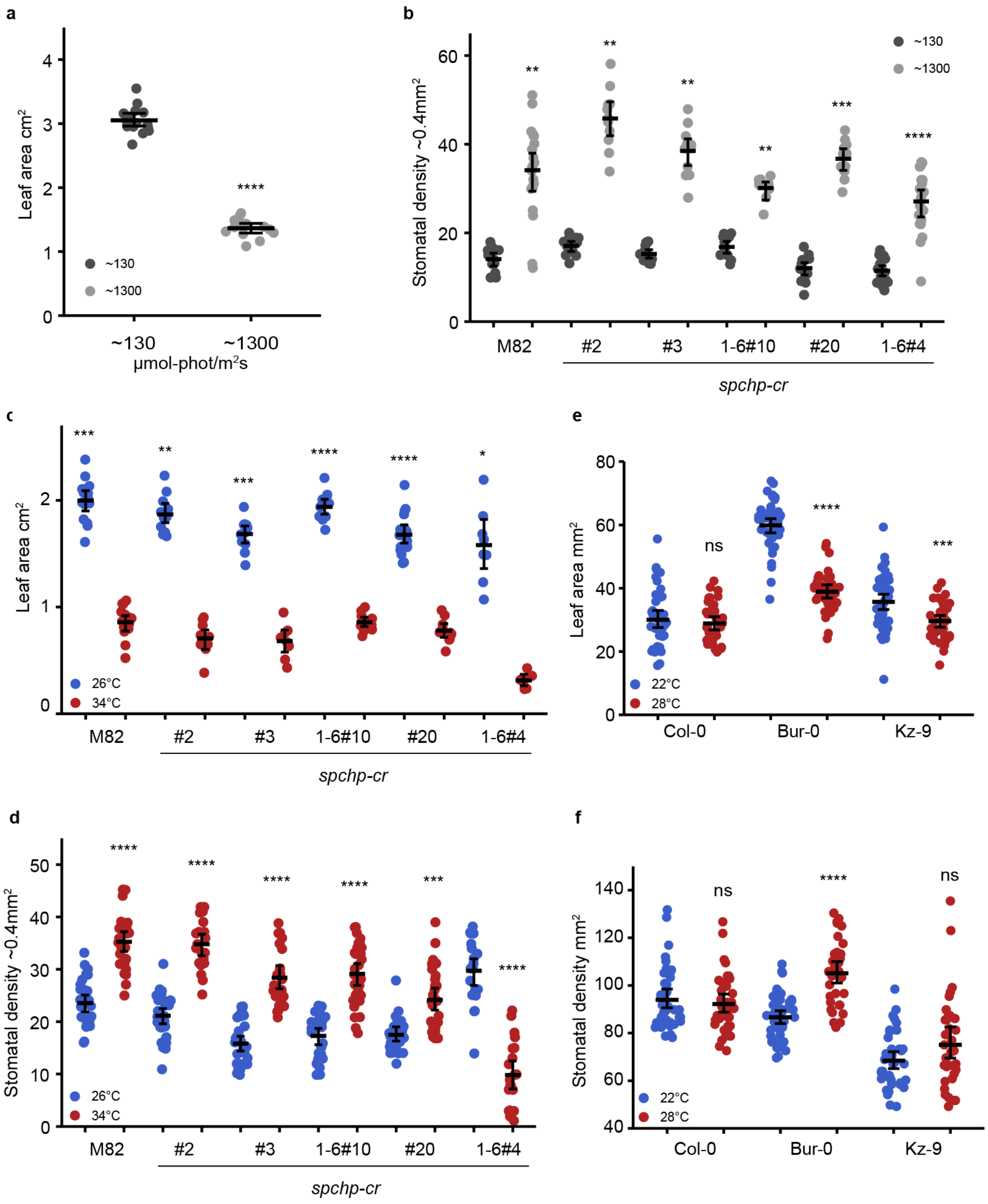


**Figure S2 Additional SI, SD and leaf area measurements of tomato and Arabidopsis variants in response to changing environmental conditions (related to Figures 2 and 3)**
(a) Plot of leaf area changes in M82 in response to light. n = 13 leaves. (b) Plot of stomatal density changes in M82 and mutants in response to light. Note the data in this plot are identical to those in Figure 2b, they have been repeated here in slightly modified form for ease of comparison. (c) Plot of in M82 and SlSPCH mutant leaf area changes in response to temperature. (d) Plot of stomatal density changes in M82 and SlSPCH mutants in response to temperature. (e) Plot of leaf area changes in Arabidopsis accessions in response to temperature. (f) Plot of stomatal density changes in Arabidopsis accessions in response to temperature. Statistical tests in (a) and (b) are represented as mean ± 95% confidence interval. Bonferroni-corrected p values from Mann-Whitney U test are *P < 0.05; **P < 0.01; ***P < 0.001; ****P < 0.0001. n.s.: P > 0.05, not significant. Sample sizes in (a) n = 13 leaves., in (b) n = 8-21 fields from 3-5 cotyledons, in (C) n = 8-18 leaves, in (d) n = 20-30 0.4 mm² fields from 4-6 cotyledons, for (e) and (f), each point represents an individual plant (n= >34).

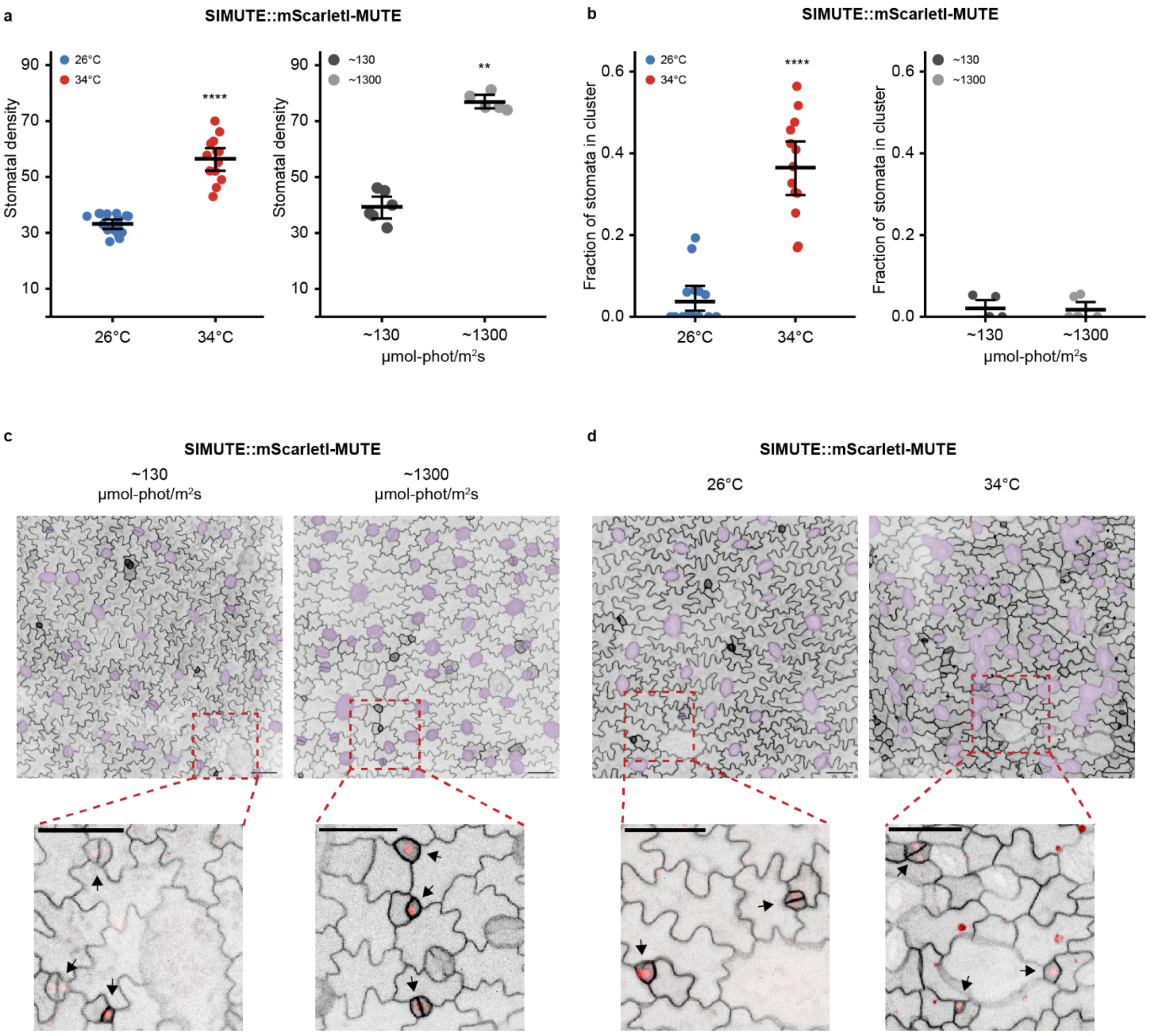


**Figure S3 Tomato lines bearing a SlMUTE translational reporter display a hypersensitive response to increased temperature, but not light (related to Figure 3)**

(a) Plots of stomatal density found in lines bearing SlMUTEpro:mScarletI-SlMUTE (hereafter, SlMUTE) in response to changes in light and temperature (b) Plots of fraction of stomata found in clusters in lines bearing SlMUTE, in response to changes in light and temperature. (c) Confocal images of 4-dpg cotyledons expressing SlMUTE at low and high light conditions. Stomata are false-colored purple. (d) Confocal images of 4-dpg cotyledons expressing SlMUTE at low and high temperature conditions. Stomata are false colored purple. Insets highlight clustered distribution of stomata consistent with "asymmetric divisions" incorrectly producing two stomata, and precocious SlMUTE expression at 34°C. Black arrows in insets point to cells expressing SlMUTE.

Scale bars in c and d represent 50 µm.Statistical tests in (a) and (b) are represented as mean ± 95% confidence interval. Bonferroni-corrected p values from Mann-Whitney U test are **P < 0.01; ****P < 0.0001. n.s.: P > 0.05, not significant. E. Sample sizes in (a) and (b) n = 5-15 0.4 $mm^2$ fields from 3 cotyledons

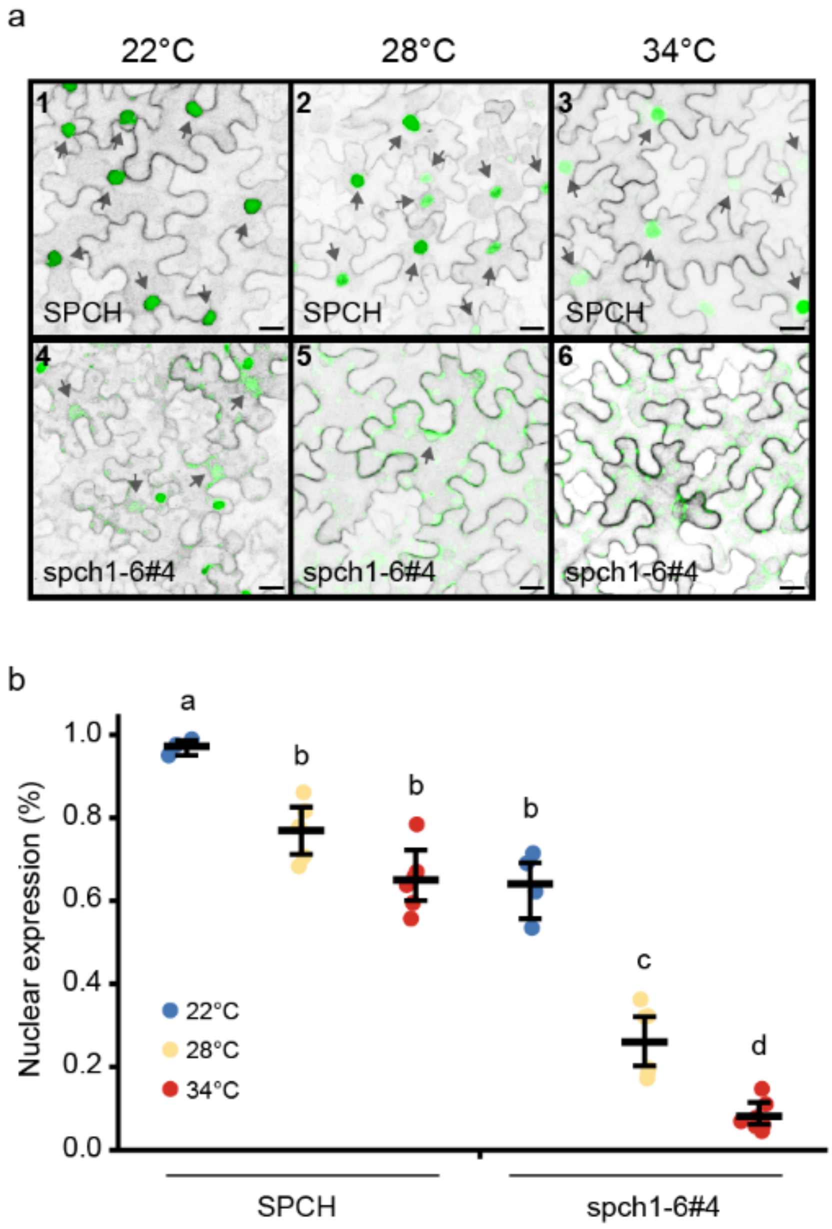


**Figure S4 Nuclear localization changes in response to high temperature in *SlSPSCH*1-6#4 (related to Figure 4)** (a) Confocal images of *N. benthamiana* leaves expressing SlSPCH full length or 1-6#4 variant as fluorescent protein fusions. As temperature rises, fewer transformed cells (confirmed by expression of PM marker) show nuclear localization of the SlSPCH 1-6#4 variant. At 34°C, full length SlSPCH also is less often present in the nucleus. Scale bars represent 20µm. (b) Quantification of data depicted in (a), with percent of transformed (PM-marker expressing) *N. benthamiana* epidermal cells that express nuclear SlSPCH variants as function of temperature. Statistical tests in (b) represented as mean ± 95% confidence interval, each letter indicates a group of samples with no statistically significant difference between their means (Bonferroni corrected p value > 0.05). Sample sizes in (a) n = 3-7 0.4 $mm^2$ fields from 3 cotyledons.

### Table S1 List of primers used for constructing reporters and for genotyping

| Primer or gRNA ID | Sequence 5′-3′ | Purpose |
|---|---|---|
| GGAG-SPCHp F | AAGAAGACTTGGAGTTCAACTTATAAAACTTTGGAATGGAGG | Golden Gate L0 promoter cloning |
| SPCHp-AATG R | AAGAAGACTTCATTTTTCAACGTTGAAAAAAGTAGAGGT | Golden Gate L0 promoter cloning |
| TTCG-SPCH F | AAGAAGACTTTTCGATGGATGGTGACCAAAATTTATCTG | Golden Gate L0 gene cloning |
| SPCH-mid F | TTGAAGACTTCCGTCTCTCCACGTATTCCTGGC | Golden Gate L0 gene cloning |
| SPCH-mid R | TTGAAGACTTACGGTtTTCAATATGACATTGGCGCC | Golden Gate L0 gene cloning |
| SPCH-GCTT R | TTGAAGACAAAAGCTTAGCAGAATGTCTGCTGAATCTGAT | Golden Gate L0 gene cloning |
| GGAG-MUTEp F | AAGAAGACTTGGAGTCTTAACCGTTTACAACTGAAAAGAATAAG | Golden Gate L0 promoter cloning |
| MUTEp-mid F | TTGAAGACTTAACGTcAGACCTATAATATATATTGTATATTGTCCAC | Golden Gate L0 promoter cloning |
| MUTEp-mid R | TTGAAGACTTCGTTTATTACCATATGGACATTTACGTT | Golden Gate L0 promoter cloning |
| MUTEp-AATG R | AAGAAGACTTCATTGATACTTTTTTTTTTCTTCTTAAAAAAAAAAAAC CTAAAT | Golden Gate L0 promoter cloning |
| TTCG-MUTE F | AAGAAGACTTTTCGATGTCTCACATAGCAGTGGAGAGAAACAGGAGgA GA | Golden Gate L0 gene cloning |

| | | |
|---|---|---|
| MUTE-GCTT R | TTGAAGACAAAAGCCTATATCTCGTTGATACATAAAACATCAGATGAGGTGAA | Golden Gate L0 gene cloning |
| GGAG-FAMAp F | AAGAAGACTTGGAGATATTTGTTTCCTAGATATTTTTTTCCATCTCACA | Golden Gate L0 promoter cloning |
| FAMAp-AATG R | AAGAAGACTTCATTTGTCTTGTTATAGTTTTTTTTTCTTTCTTTTTGTTTG | Golden Gate L0 promoter cloning |
| TTCG-FAMA F | AAGAAGACTTTTCGATGGAGAAAGAAGAAAATTGCCAGG | Golden Gate L0 gene cloning |
| FAMA-mid F | TTGAAGACTTTCATGCCTGGCTCCTATGTTC | Golden Gate L0 gene cloning |
| FAMA-mid R | TTGAAGACTTATGAGgGACCTCAATACACGAAGAT | Golden Gate L0 gene cloning |
| FAMA-C F | TTGAAGACTTGACCAGGACAACTCATAAAGGC | Golden Gate L0 gene cloning |
| FAMA-C R | TTGAAGACTTGGTCTcCTTCTTGATAGAATCTTGATCAT | Golden Gate L0 gene cloning |
| FAMA-GCTT R | TTGAAGACAAAAGCCTAGTTGATATCATATGGGGCTATTTCAGC | Golden Gate L0 gene cloning |

**Table S2 List of SlSPCH promoter sgRNAs used for CRISPR/Cas9 mutagenesis PCR primers used for genotyping**

| **Primer or gRNA ID** | **Sequence 5′-3′** | **Purpose** |
|---|---|---|
| gRNA1 | TTCAACGTTGAAAAAAGTAG | Golden Gate cloning |
| gRNA2 | TCTGCACTTTTACTTTTGTT | Golden Gate cloning |
| gRNA3 | ATATATGCCATTTTGTTGAG | Golden Gate cloning |
| gRNA4 | TTATGTTAGCAAGTTCAAAC | Golden Gate cloning |
| gRNA5 | TTTCAAATAGAAAGATACGA | Golden Gate cloning |
| gRNA6 | ATATAATCTACACATTAACA | Golden Gate cloning |
| gRNA7 | ATACTCATACTTCATAGTTA | Golden Gate cloning |
| gRNA8 | GTTTGGCTCTGCTTGAAAGC | Golden Gate cloning |
| gRNA9 | ACTTAAAAAGAAACGCGACG | Golden Gate cloning |
| gRNA10 | ATAAATATGATATATTATGG | Golden Gate cloning |
| gRNA11 | GGGAATAAGAGAGAGAGCAA | Golden Gate cloning |
| SlSPCHpro F | TCTCGTTTACGGAAAGTAGAAACC | Genotyping, cloning and sequencing |
| SPCH155 R | ATTGTTGCAGTCTGGTTGTTGAT | Genotyping, cloning and sequencing |
| SlSPCH2937up F | GCGTCGTCGGAAAATCCAAG | Promoter sequencing |
| SlSPCH2168up F | GAGCTTGTTTGGCTCTGCTTG | Promoter sequencing |
| SlSPCH1700up F | TGCTACATTAATTACATTGAGTGAGAA | Promoter sequencing |
| SlSPCH1311up F | CAAATAGAAAGATACGAAGGTCGAAGA | Promoter sequencing |
| SlSPCH943up F | AAGAACCCGTTTGAACTTGC | Promoter sequencing |
| SlSPCH F | TATTCCCTTTTCTACATCTCCTCTTT | Genotyping, cloning and sequencing |

| | | |
|---|---|---|
| SlSPCH_R | TTCTCTTATTATTGTTACAGCTCCCTA | Genotyping, cloning and sequencing |
| SlSPCH200 | GGTGTCAGAAGAAGGAAAGAAGA | Gene sequencing |
| SlSPCH386 | ACTTTGATGCCTTGTTTTTATGCTAA | Gene sequencing |
| SlSPCH951 | GGTGATCAAGCATCAATAATTGGT | Gene sequencing |
| SPCH1371 | CCGTCTCTCCACGTATTCCTGGC | Gene sequencing |
